# Tropism, susceptibility, infectivity, and cytokine releases of differentiated human tonsillar epithelial cells by different Influenza viruses

**DOI:** 10.1101/2021.05.03.442542

**Authors:** Faten Okda, Ahmed Sakr, Robert Webster, Richard Webby

**Affiliations:** Department of Infectious Diseases, St. Jude Children’s Research Hospital, Memphis, Tennessee, USA; Department of business Administration, Dakota state University, Madison, USA

**Author notes:** Address correspondence to Richard Webby,.

**Keywords:** Tonsils, crypts, squamous epithelial cells, keratin, reticular epithelial cells, adaptation, pathogenesis, cytokines, chemokines, influenza virus, H1N1, H3N2, WSN

## Abstract

Human tonsil epithelium cells (HTEC) are a heterogeneous group of actively differentiating cells comprising stratified squamous epithelial and reticulated crypt cells with abundant keratin expression. We hypothesized that the tonsils are a primary site for influenza infection and sustained viral replication. Primary HTEC were grown using an air-liquid culture and infected apically with different influenza viruses (IVs) to measure viral growth kinetics. These cultures were highly differentiated, with subpopulations of heterogenous surface stratified squamous cells rich with both cilia and microvilli; these cells contained more α2,6-linked sialic acids, those preferentially bound by human IVs, than α2,3-linked avian like sialic acids. The stratified squamous cells were interrupted by patches of reticular epithelial cells rich in α2,3-linked sialic acids. The HTEC were permissive for influenza A and B virus replication. Following infection, a subset of cells, mostly ciliated cells, underwent apoptosis while others remained intact despite being positive for IV nucleoprotein. H3N2 virus antigen colocalized with non-ciliated cells while H1N1 virus antigen was mostly associated with ciliated cells. Exposure of HTECs to IVs triggers an early proinflammatory response that fluctuates between viruses. The H3N2 IV induces an early response that persists, whereas pH1N1 induces a primarily late response in HTECs. Our results implicated HTEC as a site for IV replication. The HTEC differentiated system provides a valuable in vitro model for studying cellular tropism, infectivity, cytokine responses and the pathogenesis of IVs.

**IMPORTANCE:** To develop an effective intervention against influenza, it is important to identify host factors affecting transmission, pathogenesis, and immune response. Tonsils are lymphoepithelial organs characterized by infiltration of B and T lymphocytes into the squamous epithelium of tonsillar crypts, beneath which germinal centers play key roles in antigen processing and immune response. The heterogenicity of HTECs as well as the sialic acid distributions supports the replication of IVs and may play a role in IV adaptation. Furthermore, Tonsillectomy is a surgical procedure in which tonsils are fully removed from the human throat and may contribute to the diverse outcomes among infected individuals.

## INTRODUCTION

Influenza viruses (IVs) are an important health threat due to annual disease outbreaks and pandemic risks (1–3). To date, no effective universal vaccine against IVs has been developed partly because of a limited understanding of IV pathogenesis (4). Identifying the host and virologic factors contributing to IV disease severity are critical to influenza control (1, 4). The tonsils are lymphoid organs located on either side of the back of the human throat (5) that consist of heterogeneous lymphoepithelial cells. The tonsils are comprised of two different types of actively differentiating epithelia: the lining stratified squamous epithelium and the reticulated crypt, or lymphoepithelium (6).

Each human tonsil contains a network of pits known as crypts, which are branched invaginations lined by stratified squamous epithelia that extend throughout the full thickness of the tonsil. The tonsillar crypts represent a specialized compartment that is critical for the immunologic functions of the tonsils. The tonsillar crypts play a role in facilitating contact between environmental factors and lymphoid tissues, in which many lymphoid cells pass through. These lymphoid cells may escape into the epithelium and mix with the saliva to form salivary corpuscles (7).

The surface stratified squamous epithelium differs from a simple ciliated epithelium because it acts as an extracellular matrix and surface, controls tissue integrity, and mediates intracellular signaling pathways and immunologic functions (6, 8). The tonsil epithelium monitors and protects the body against respiratory and gastrointestinal infections (9). Typically, lymphocytes infiltrate into the squamous epithelium of the tonsillar crypts and mucosal surface. The tonsils have several local and systemic immunologic functions that involve innate, cellular, and humoral immunity (6). The tonsils confer immunity against entering pathogens and enable efficient and rapid systemic immune responses by antigen sampling, intracellular signaling pathways, and movement of lymphocytes, cytokines, and chemotactic molecules from the tonsils to other lymphoid organs. The tonsils contain a germinal center, in which B memory cells and secretory antibodies (IgA) are produced. Activated cells in mucosa-associated lymphoid tissues primarily secrete local IgA-type antibodies (6, 9). However, the role of the tonsil epithelia in IV pathogenesis remains uncertain.

Children are susceptible to tonsillitis because their tonsils are more active and have been exposed to fewer pathogens than those of adults. Tonsillectomy procedures fully remove the tonsils from the human throat (10). Tonsillectomy data collected over the last 60 years reveal decreased trends of tonsillectomy procedures performed in the United States (11, 12); however, the percentage of human subpopulations without tonsils is high at different ages (12, 13). Tonsil removal is considered a controversial environmental factor in inflammatory diseases, such as Crohn disease, because of the depletion of CD4^+^/CD25^+^ T-regulatory cells (14). Tonsillectomy is also associated with an increased risk of Behcet disease (15), poliomyelitis (16–18), and respiratory, infectious, and allergic diseases (19). Of 1.2 million Danish children who received tonsillectomies, 43 207 had increased risk of developing respiratory, infectious, and allergic conditions. High mortality risk was reported in young adults who received tonsillectomies because of poor lung function (20). Moreover, an increased risk of autoimmune conditions such as thyroid disease, rheumatic diseases, and type 1 diabetes occurred in 179 875 Swedish patients who received tonsillectomies (21). Tonsillectomy is also associated with a higher risk of cancer (22–24) and premature acute myocardial infarctions (25). The prevalence of IV infections, disease severity, and immunity levels in patients without tonsils is poorly defined.

The tonsils are an initial site of replication of foot-and-mouth disease virus (FMDV) (26), polyomavirus (27), HIV (28), and Epstein–Barr virus (5, 8). Porcine tonsils are primary infection sites for FMDV after simulated natural virus exposure and facilitate replication in the clinical phase, yielding high amounts of viral shedding into the environment due to extensive amplification (26). HIV specifically targets the tonsils because of the interaction with HIV targets in this tissue (28).

We hypothesize that the tonsils are a primary site for IV infection and contribute to viral pathogenesis. IVs cause febrile respiratory infections ranging from self-limited to severe symptoms in children and adults. Children aged 5–14 years with pandemic H1N1 (pH1N1) infections experienced a high rate of hospitalizations, including acute respiratory distress syndrome (29, 30). The virulence of pH1N1 in children is linked to the downregulation of type 1 interferon expression, apoptosis, and hyperinduction of proinflammatory cytokines (31). In contrast, the 1918 H1N1 pandemic IV caused severe infections in healthy individuals aged 15 to 34 years (32, 33) because of exaggerated proinflammatory immune responses (34). Upregulation of inflammatory cytokines and chemokines are associated with the severity of influenza (35), which may benefit from immunomodulatory therapy. Identifying the pathways involved is critical. Here, we investigated the pathogenesis of IVs in a human tonsillar epithelial cell (HTEC) culture model to glean insight into the contribution of the tonsils to the diverse outcomes and immune responses of IV infections in humans.

## RESULTS

### Tonsillectomy

We analyzed the number of tonsillectomies performed within the last 40 years in the United States. A high percentage of tonsillectomies have been performed, leading to 25% to 60% of the US population without tonsils (Fig. 1).

**FIG 1.**
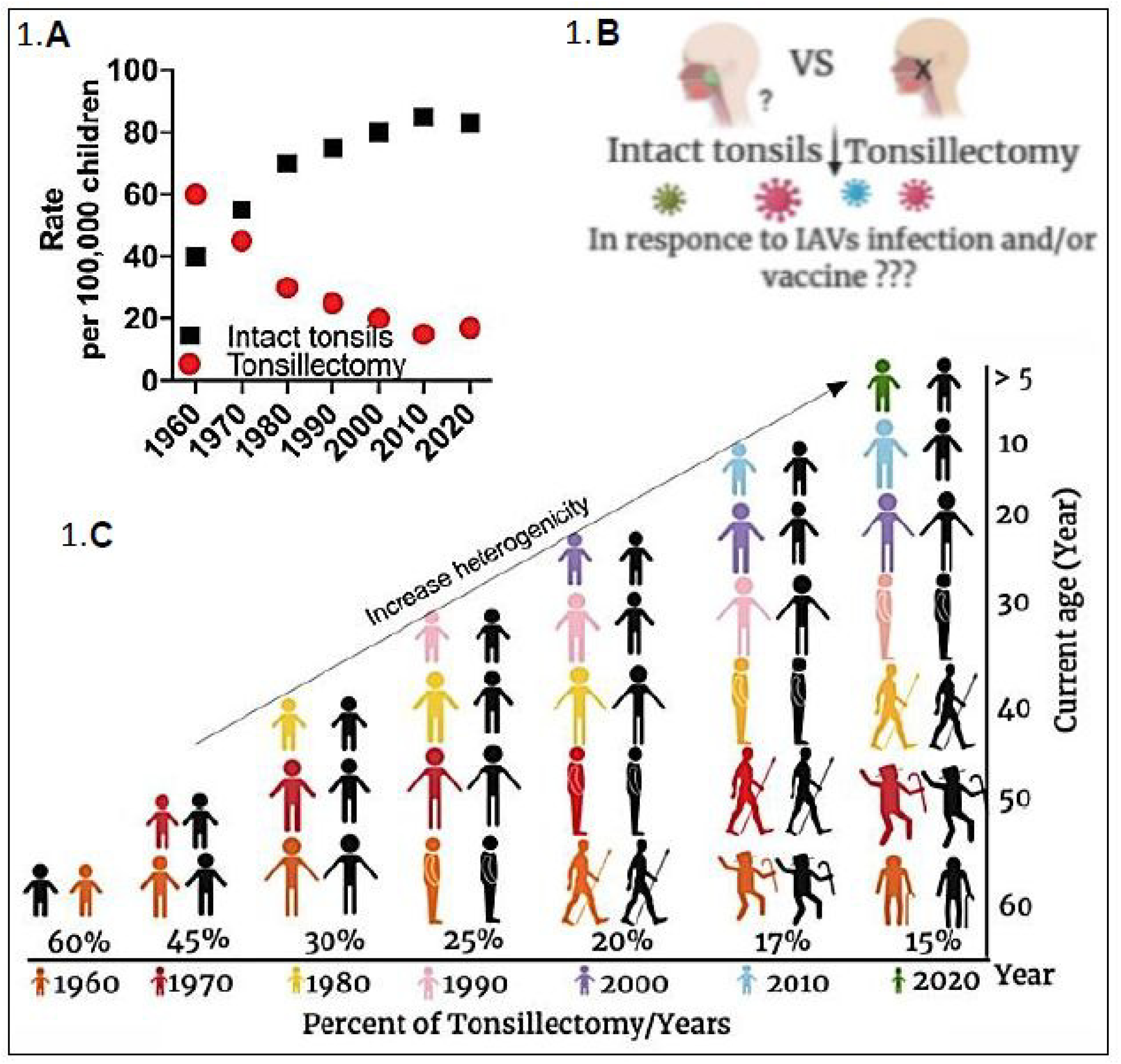
Tonsillectomy trends in the United States and heterogenicity by age. (A) Trends of patients receiving tonsillectomies (red) *vs.* those who did not (black) in the United States from 1960 to 2020. (B) The tonsils are located in the nasopharynx of humans. (C) Graphical illustration of the percentage of tonsillectomies performed per year and increased heterogenicity by age. Each color represents the decade of childhood tonsillectomy and subsequent growth until 2020. Black represents the populations with intact tonsils.

### Human tonsillar epithelial cell culture model

To elucidate the contribution of the tonsils to IV pathogenesis and immune responses, we established a highly differentiated HTEC model. Phase-contrast microscopy images revealed that the HTECs were heterogeneous, with different cell sizes, types, and cytoplasm-to-nucleus ratios. Some HTECs acquired a very large cytoplasm-to-nucleus ratio, indicative of differentiated squamous epithelial cells, and subsets of differentiated morphologically large epithelial cells appeared in the cultures over time (Fig. 2A). Scanning electron microscopy (SEM) imaging revealed that the apical surface of the HTECs contained microvilli of approximately 5 *μ*m, and few ciliated epithelial cells were present (Fig. 2B).

**FIG 2.**
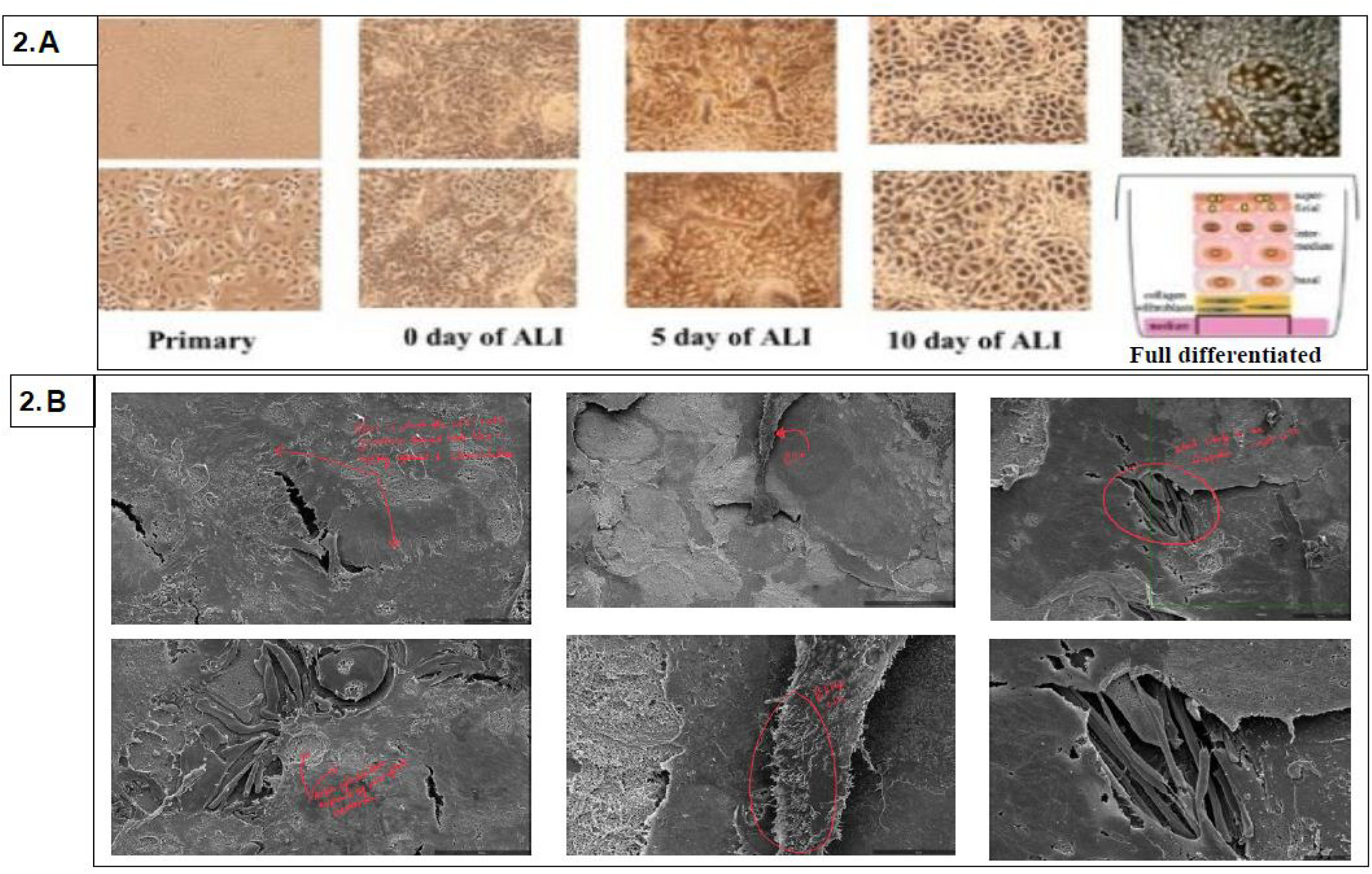
Characterization of well-differentiated human tonsil epithelial cells. (A) Phase contrast microscopy images of day 10 human tonsillar epithelial cell (HTEC) cultures at the air–liquid interface reveals well-differentiated epithelial cells of different sizes, types, and cytoplasm-to-nucleus ratios. (B) Scanning electron micrograph illustrating the apical surface of the HTECs, which contain few ciliated epithelial cells and more microvilli. Scale bars = 5 *μ*m.

### Morphological examination of well differentiated HTEC cultures

To further characterize our HTEC model, we stained the cells with a panel of monoclonal antibodies against epithelial K5 and K14 as markers of surface stratified epithelial cells (Fig. 3A) and K8/18 and K19 (Fig. 3B) as markers of the tonsillar crypts. We also stained the HTECs with antibodies against *β*-tubulin as a marker of cilia and villin as a marker of microvilli. We found that the differentiated HTECs consisted of a layered surface stratified epithelium with K5^+^ and K14^+^ cells (Fig. 3A). Although the K5^+^ cells exhibited abundant cilia and microvilli, the K14^+^ cells had more microvilli and less cilia (Fig. 3A). The tonsillar crypt structures detected by K8/18 and K19 expression had an abundant distribution of cilia and microvilli but more microvilli and fewer cilia (Fig. 3B). Infiltration of the intercellular spaces by lymphocytes was also evident in the HTECs (Fig. 3).

**FIG 3.**
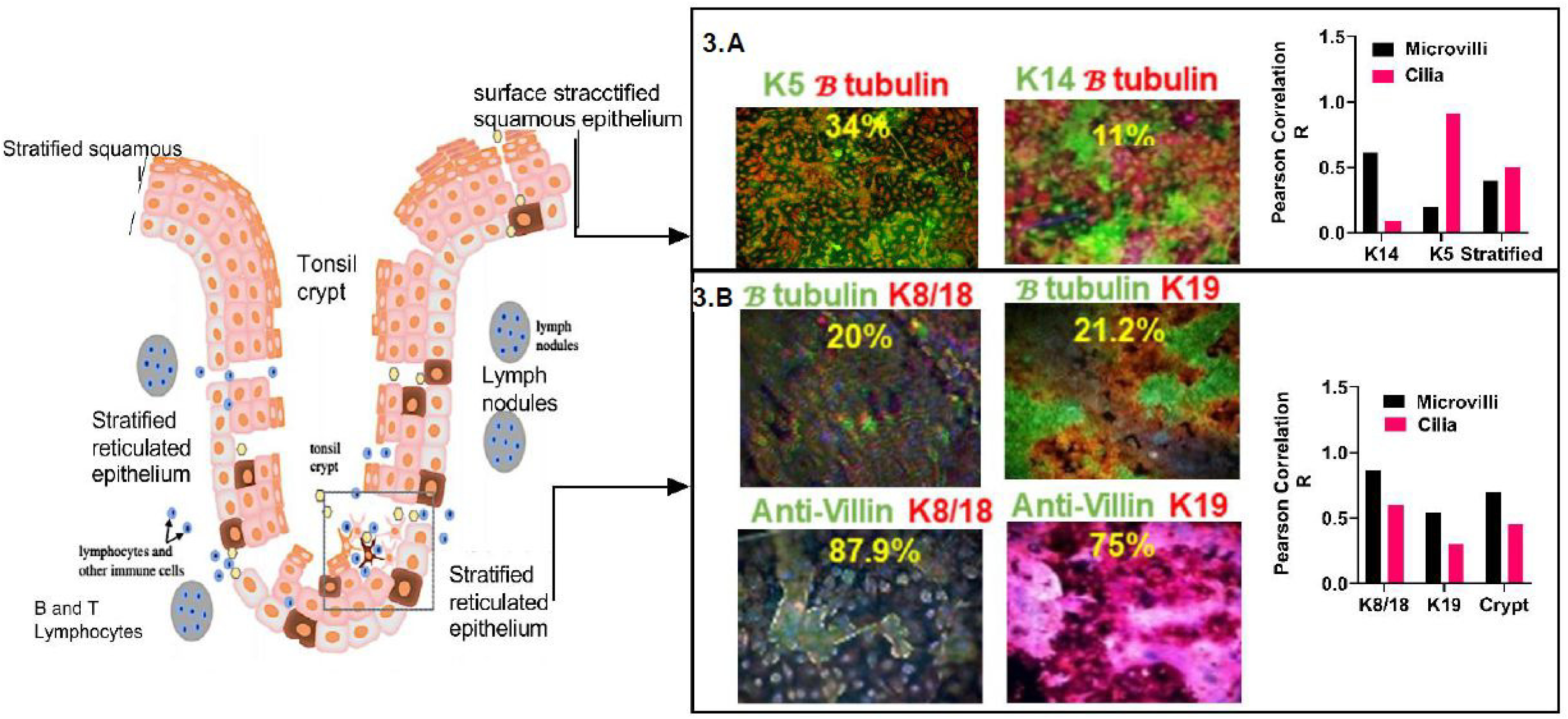
Structure of the tonsil tissue and keratin expression in HTECs. A schematic of the tonsil structure in the left side adapted from (74) and has been modified. In the right side (A) Confocal microscopy showing that the tonsillar extracellular epithelial surface (K5/K14^+^) contains few ciliated cells (*β*-tubulin^+^) and (B) that the crypts (K8/18^+^ and K19^+^) have more villi and microvilli than cilia. Cells were fixed with chilled acetone and stained with monoclonal antibodies against K5, K14, K18, K19, *β*-tubulin, and villin. Colocalization was determined by using ImageJ and calculated by the Pearson correlation (*R*).

### Sialic acid distribution in HTECs

To determine the sialic acid (SA) distribution in the well-differentiated HTECs, we used the *Sambucus nigra* agglutinin (SNA) and *Maackia amurensis* agglutinin II (MAA II) lectins to visualize avian- *α*2,6- and human-virus *α*2,3-linked SA receptors, respectively. Both *α*2,3 and *α*2,6-linked SA receptors (green) were present on both the surface stratified (K5^+^ and K14^+^, red) and crypt epithelia (K8/18^+^ and K19^+^, red) (Fig. 4). Few ciliated cells and surface stratified epithelial cells were rich in human-virus *α*2,6-linked SA receptor (Fig. 4A, C). However, the microvilli-containing epithelial cells and reticular crypt cells were rich in *α*2,3–linked SA (Fig. 4B, C).

**FIG 4.**
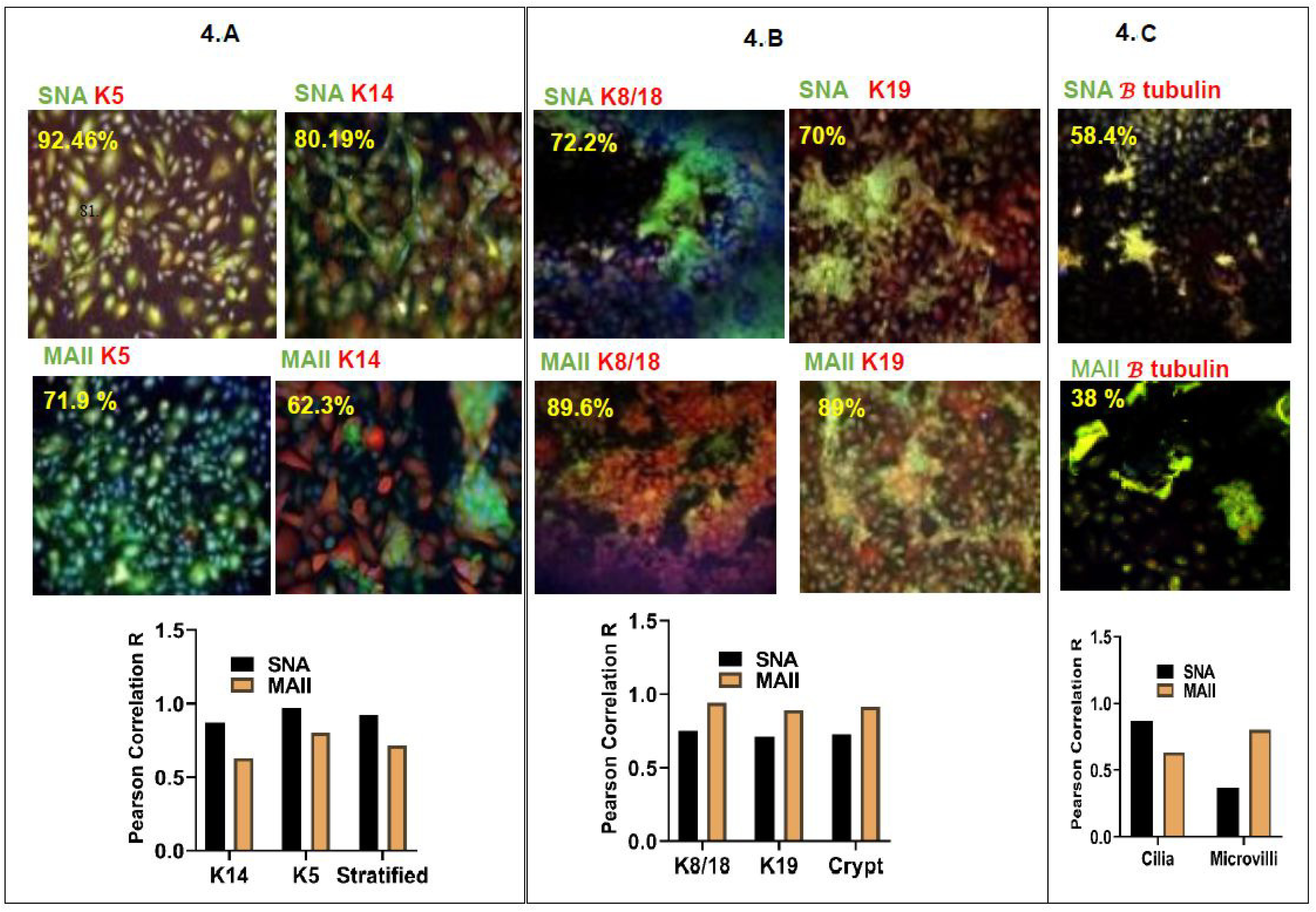
Lectin and keratin staining of human tonsillar epithelial cells. (A) The surface stratified epithelium (K5/K14^+^) is richer in *α*2,6-linked SA receptors (SNA^+^) than in avian-like *α*2,3-linked SA receptors (MAII^+^). (B) The reticular crypts (K19^+^ and K8/18^+^) are richer in *α*2,3-linked SA receptors (MAII^+^) than in *α*2,6-linked SA receptors (SNA^+^). (C) Lectin staining of the distributed cilia. The cells were fixed with chilled acetone and stained with monoclonal antibodies against K5, K14, K18, K19, *β*-tubulin, and villin. Colocalization was determined by using ImageJ and calculated by the Pearson correlation (*R*).

### Susceptibility of HTECs to IV infection

To investigate the susceptibility of the HTECs to IV infection, we used various IVs (Table 1) to infect the HTECs *in vitro*. If the tonsillar epithelia are important for IV pathogenesis, adaptation, and transmission, these cells will be susceptible to infection with swine, avian, and human IVs. Indeed, we found that the HTECs were permissive for the growth of various A and B IVs (Table 1).

**TABLE 1.** Susceptibility of HTECs to different IVs.

| Virus | 24 h post infection |  |  | 48 h post infection |  |  |
| --- | --- | --- | --- | --- | --- | --- |
|  | HA<sup>a</sup> | MFI<sup>b</sup> | PFU<sup>c</sup> | HA<sup>a</sup> | MFI<sup>b</sup> | PFU<sup>c</sup> |
| A/SWITZERLAND/8060/2017(H3N2) | N | 3725 | $6.0 \times 10^5$ | N | 4120 | $2.1 \times 10^6$ |
| A/HONG KONG/4801/2014 (H3N2) | 64 | 17 890 | $3.0 \times 10^5$ | 64 | 20 198 | $4.7 \times 10^6$ |
| A/MEMPHIS/257/2019 (H3N2) | 64 | 20 156 | $1.2 \times 10^6$ | 128 | 23 876 | $6.8 \times 10^6$ |
| A/SWINE/OH/16TOSU4783/2016 (H3N2) | 32 | 4993.5 | $2.0 \times 10^4$ | 32 | 5321 | $4.7 \times 10^4$ |
| A/TENNESSEE/1-560/2009 (H1N1) | 256 | 18 770 | $3.4 \times 10^7$ | 256 | 19 064 | $4.2 \times 10^7$ |
| A/BRISBANE/59/2007 (H1N1) | 64 | 22 134 | $2.0 \times 10^6$ | 64 | 21 350 | $3.0 \times 10^6$ |
| A/SWINE/MN/37866/1999 (H1N1) | 64 | 20 882 | $1.5 \times 10^5$ | 32 | 21 071 | $2.6 \times 10^5$ |
| A/SWINE/NC/18161/2002 (H1N1) | 32 | 5445. | $2.0 \times 10^4$ | 32 | 5046 | $1.2 \times 10^5$ |
| A/SWINE/OH/09SW1486/2009 (H1N2) | 16 | 5235 | $3.5 \times 10^4$ | 32 | 5650 | $7.0 \times 10^5$ |
| RGA/VIETNAM/1203/04 (H5N1) | N | 6900 | $1.3 \times 10^5$ | N | 6440 | $4.1 \times 10^5$ |
| C/JOHANNESBURG/1/1966 | N | 450 | —<sup>d</sup> | N | 1072 |  |
| B/BRISBANE/60/08 (EGG) | 64 | 3704 | $5.6 \times 10^5$ | 64 | 4250 | $4.2 \times 10^5$ |
| B/BRISBANE/60/08 (CELL) | 32 | 2225.5 | $3.0 \times 10^5$ | 32 | 3420 | $2.5 \times 10^4$ |
| A/RG/WSN/33 | 64 | 191340 | $2.7 \times 10^7$ | 128 | 23 498 | $7.3 \times 10^7$ |

<sup>a</sup>HA, hemagglutinin assay. <sup>b</sup>MFI, mean fluorescence intensity. <sup>c</sup>PFU, plaque-forming unit. <sup>d</sup>The symbol “—” means undetermined.

### Viral replication kinetics

To analyze the replication of WSN, pH1N1, and seasonal H3N2 IVs in HTECs, we infected them from the apical side of the air–liquid interface. We analyzed single-step (Fig. 5A) and multistep growth curves (Fig. 5B). With single-step replication, we found that the titers of released WSN were markedly higher than those of pH1N1 and H3N2 after 4 h of infection (Fig. 5A). WSN titers began to decline at 12 h PI (post infection) at a point where H3N2 titers were highest. The H3N2 IV maintained high titers until 40 h post infection, whereas pH1N1 exhibited late replication kinetics, with high viral titers at 24 h PI (Fig. 5A). To understand the long-term influence of IV infection in HTECs, we analyzed multiple-step kinetics of the three IVs. The H3N2 and WSN IVs exhibited high viral titers from day 1 PI and were maintained until day 8 post infection. In contrast, pH1N1 grew slower, and release began at day 3 PI (Fig. 5B).

**FIG 5.**
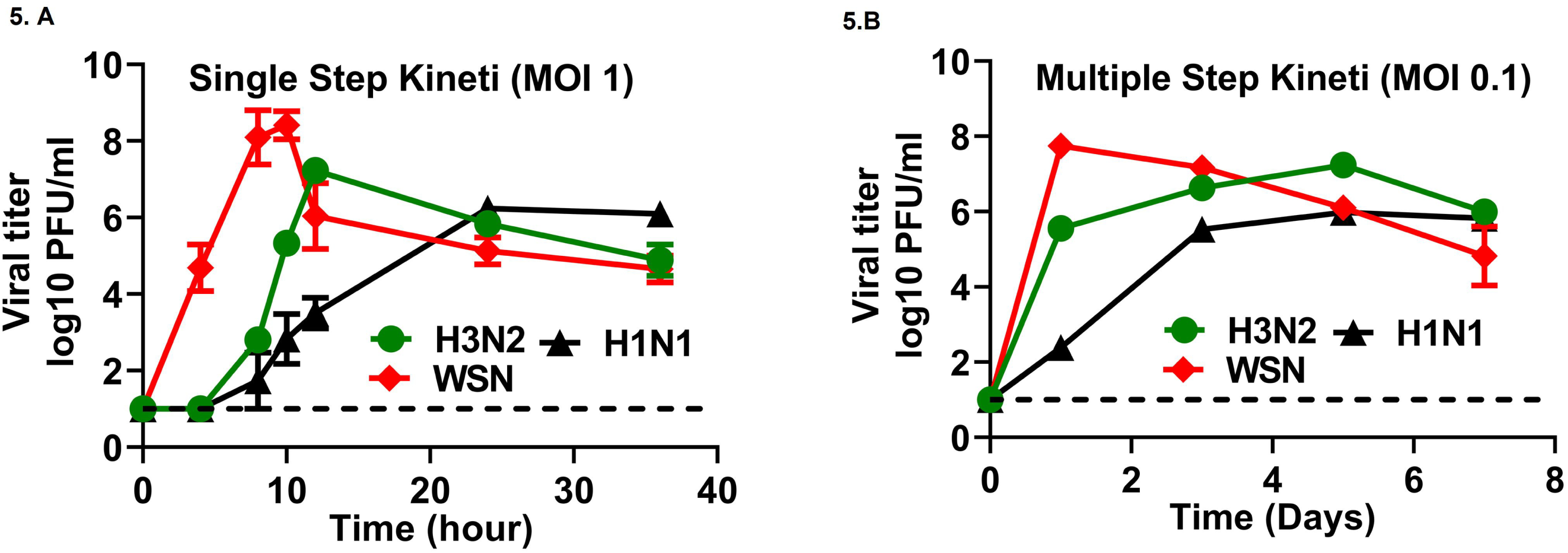
(A) Single-step H1N1, H3N2, and WSN (MOI 1) replication kinetics in HTECs. (B) Multiple-step H1N1, H3N2, and WSN (MOI 0.1) replication kinetics in HTECs. Well-differentiated HTECs were inoculated with IVs on the apical side. IVs released from the apical side were harvested at different time points and titrated by plaque assay. The means ± SEM of five HTEC cultures from three independent experiments are shown. Statistical significance was determined by two-way ANOVA and Tukey posthoc analysis, comparing each set of results to those for the mock cells. \**P* < 0.05; \*\**P* < 0.01; \*\*\**P* < 0.001; \*\*\*\**P* < 0.0001.

### Colocalization of IVs subtypes with well-differentiated HTEC

To understand the long-term effects of IV infection in human tonsil cells, we performed quantitative analyses of the levels of infected HTEC surface stratified epithelial and crypt cells, as well as those characterized by cilia and microvilli. We infected well-differentiated HTECs with pH1N1 or H3N2 from the apical surface at a multiplicity of infection (MOI) of 0.1 and stained the cells at days 2, 5, and 7 PI to detect viral nucleoprotein (red), K5 (green), K14 (green), K8/18 (green), K19 (green), and cilia (green). The H3N2 IV was highly associated with reticulated noncilited cells during initial replication (Fig. 6A). At day 2 post infection, the surface stratified epithelial and crypt cells were mildly infected with H3N2. At day 5 post infection, both the surface stratified and crypt cells were infected. At day 7 post infection, the crypt cells were mildly infected. Loss of surface stratified epithelial cells occurred in all virus infected HTECs, as compared with that of mock cultures at day 7 post infection.

**FIG 6.**
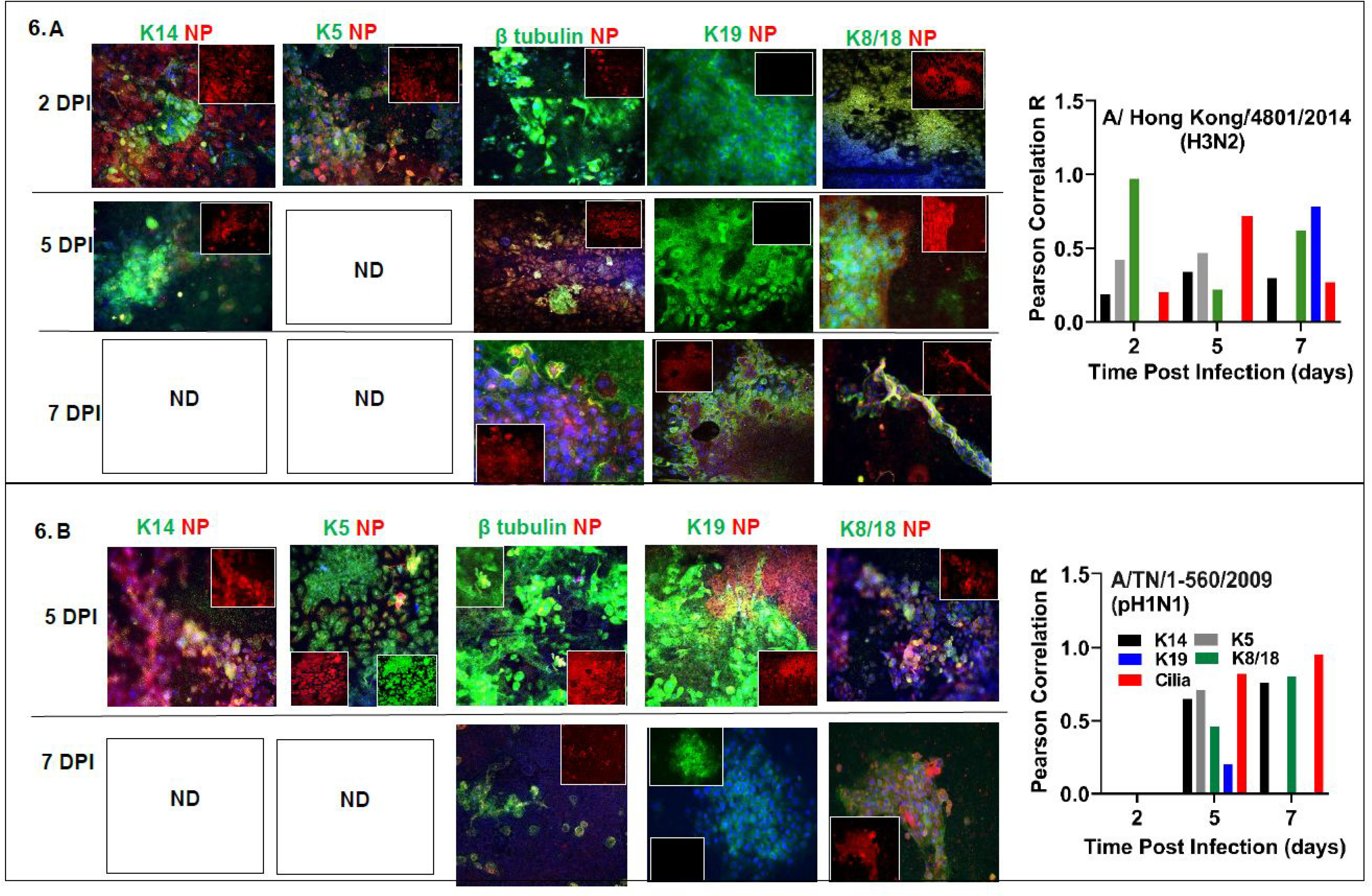
Colocalization of IV nucleoprotein staining in HTECs. (A) H3N2 colocalized more with the tonsillar crypts than with surfaces. (B) H1N1 associated more with the ciliated and surface stratified epithelia than with the crypts. The crypts remained intact despite infection. Well-differentiated HTECs were infected with IVs on the apical side at a MOI of 0.1. Epithelial cells were fixed with chilled acetone at days 2, 5, 7, and 10 PI and stained with monoclonal antibodies against K5, K14, K18, K19, *β*-tubulin, and IV nucleoprotein. Colocalization was determined by using ImageJ and calculated by the Pearson correlation (*R*).

The pH1N1 IV primarily infected ciliated cells and replicated efficiently in the surface stratified epithelium, despite a low and delayed initial infection rate (Fig. 6B). At day 2 post infection, no viral nucleoprotein antigen was detected. However, at day 5 post infection, both the surface stratified and crypt epithelial cells were infected, with abundant IV present in the ciliated cells. Of note, the number of ciliated cells was clearly increased at this time. At day 7 post infection, the surface stratified epithelial cells were lysed, and the crypt cells remained intact, with infection of some ciliated cells but less than at 5 days post infection. The microvilli were infected with all of the IVs but colocalized more with the H3N2 virus. A subset of infected cells with pH1N1, mostly ciliated, underwent apoptosis, whereas others (including non-ciliated cells) remained intact despite IV nucleoprotein positivity. At day 2 post infection, all of the cell types were infected with the WSN virus and the surface stratified epithelial cells were lysed. The WSN virus replicated efficiently in both the surface stratified epithelial and crypt cells.

These results demonstrate that the H3N2 virus preferentially infects microvilli-rich cells, replicates early in surface stratified epithelial cells, and later infects the crypt cells, which remain intact. The pH1N1 IV preferentially infected ciliated cells and some non-ciliated cells with delayed infection. This indicates that the heterogenicity of HTECs supports the replication of various IVs and may play a role in IV adaptation.

### Cytokine and chemokine responses in HTECs

Excessive cytokine responses are associated with severe influenza symptoms. To understand the cytokine profile of IV-infected HTECs, we examined the induction of various inflammatory cytokines and chemokines in response to infection with the various IV subtypes. We measured a panel of 49 cytokines in the supernatants from infected HTECs. We found that HTECs produced different chemokines and cytokines in response to pH1N1 and H3N2. The H3N2 IV upregulated VEGF, RANTES, MCP-1, IP-10, IL-8, IL-6, IL-1*α*, IL-1*β*, GRO, INF-*α*2, fractalkine, GM-CSF, TGA, FGF-2, and EGF during the course of infection. PDGF-AB/BB, IL-1β, PDGF-AA, and TNF-*α* were secreted only at days 3 to 5 post infection. In contrast, TNF-*β*, IL-5, IL-4, IL-2, IL-9, IL-13, and IL-12 were downregulated in H3N2-infected cells (Fig. 7A). Infection with pH1N1 resulted in the same pattern as that of H3N2, but PDGF-AA and TNF-*β* were upregulated during infection, and PDGF-AB/BB was downregulated (Fig. 7B). There was a significant INF*α*-2 release by the WSN-infected HTECs at 96 h PI (*P* < 0.0005, *vs* mock) and by pH1N1-infected cells at 120 h PI (*P* < 0.005, as compared with that of mock-infected cells and H3N2 *P* < 0.0.05) (Fig. 8A) with no significant release of INF-*γ* among the three viruses (Fig. 8B). HTEC infected with H3N2 showed an early, abrupt, and strong release of TNF-*α* at 24, 96, and 120 h PI (P < 0.05, < 0.005, and < 0.05, respectively), whereas pH1N1 cause delayed peak at 120 h pi (P < 0.0005, *vs.* mock-infected cells) (Fig. 8C). This indicate that HTECs infected with IVs failed to evoke an early interferon response, but the induction of TNF-*α* may play a pathologic role. IL-1R-1 exhibited delayed release in cells infected with pH1N1 at 120 h PI (*P* < 0.005), which occurred in higher concentrations than that in H3N2-infected cells earlier at 24 h (*P* < 0.05) (Fig. 8D). In contrast, the IL-1a levels released in WSN-infected cells were lower than that in H3N2-infected cells at 24 h PI (P < 0.05) (Fig. 8E). While IL-1*β* was significantly induced only by pH1N1-at 72 h PI (Fig. FI). IL-6 exhibited early release from the HTECs infected with H3N2 at 24 and 48 h PI (*P* < 0.05 for both comparisons) while pH1N1 caused significant IL-6 release at 48, 72, and 120 h PI (*P* < 0.05, 0.005, and 0.05, respectively) with minor changes by WSN, as compared with that of pH1N1-infected cells at 72 h PI (*P* < 0.05) (Fig. 8G). There were no significant changes in IL-7 levels among the 3 viruses (Fig. 8H). While, IL-15 was significantly induced in H3N2-infected cells at 96 and 120 h PI (P < 0.005 and 0.0005, respectively) and higher in pH1N1-infected cells at 120 h PI (*P* < 0.05) (Fig. 8I). The synergistic interaction between IL-6 and IL-15 suggests that IV-infected HTECs exacerbate clinical outcomes during IV infections. The H3N2 and pH1N1 infected HTECs induced IP-10 (CXCL10) at 120 h PI (*P* < 0.005 and 0.05, respectively, *vs.* mock infection), with hyperactivated proinflammatory IP-10 response by pH1N1 than H3N2 (*P* < 0.05) (Fig. 8J). Unlike the previous cytokines, G-CSF was significantly induced by pH1N1 as early as at 72 h PI (*P* < 0.05) and late by H3N2 at 120 h PI (*P* < 0.0005) (Fig. 8K). However, GM-CSF levels were higher in H3N2-infected cells than in pH1N1-infected cells (*P* < 0.05) and WSN-infected cells (*P* < 0.05) at 24 and 96 h PI (Fig 8L). The release of G-CSF and GM-CSF from the HTECs indicates that the tonsils may contribute to the immune/inflammatory cascade induced early by H3N2 and after 4 days by pH1N1. The chemokine CCL2 (MCP-1) showed early increased by H3N2 (*P* < 0.05, *vs.* mock infection, and *P* < 0.05 *vs*. pH1N1) at 24 h PI with fluctuating level at 48 and 72 h PI. While pH1N1 showed late peak at 120 h PI (Fig. 8M). There were no significant changes of either MIP1b or MIP1a levels with any of the IVs (Fig. 8N, O). While, the VEGF was upregulated by H3N2 and WSN at 24 h PI, as compared with that of mock-infected cells (*P* < 0.05 and 0.05, respectively), and higher than pH1N1 (*P* < 0.05 and 0.05, respectively) at 48 h PI (Fig. 8P). This indicates that H3N2 enhances early healing and regeneration. HTECs infected with WSN released fractalkine (CX3CL1) at 24 h PI, with maximum increase at 48 h PI, in contrast with that of mock- and H3N2-infected cells (*P* < 0.0005 and 0.005, respectively). However, H3N2-infected HTECs induced significantly higher fractalkine at 96 h PI and pH1N1 at 120 h PI (Fig. 8Q). There was a fluctuating release of the RANTES by H3N2 at 24 and 120 h PI (*P* < 0.0005 and 0.005, respectively), and late release by pH1N1 at 120 h PI (*P* < 0.05) (Fig.7R). None of the IVs induced significant levels of GRO in the HTECs (Fig.7S). PDGF-AA was induced only at 48 h PI by H3N2 (*P* < 0.05) with higher levels than that of pH1N1 (*P* < 0.05) (Fig. 8T). While FGF-2, EGF and TGF-*α* induced at 24 h (*P* < 0.05) and 120 h (*P* < 0.0005) PI by H3N2 with significant delayed exaggeration by pH1N1 at 120 h PI (*P* < 0.05) (Fig. 8U-W). Together, these findings indicate that HTECs play a distinct role in the induction of various proinflammatory and chemoattractive cytokines during IV infection which fluctuates during infection and H3N2 induces an early response that persists, whereas pH1N1 induces a primarily late response in HTECs.

**FIG 7.**
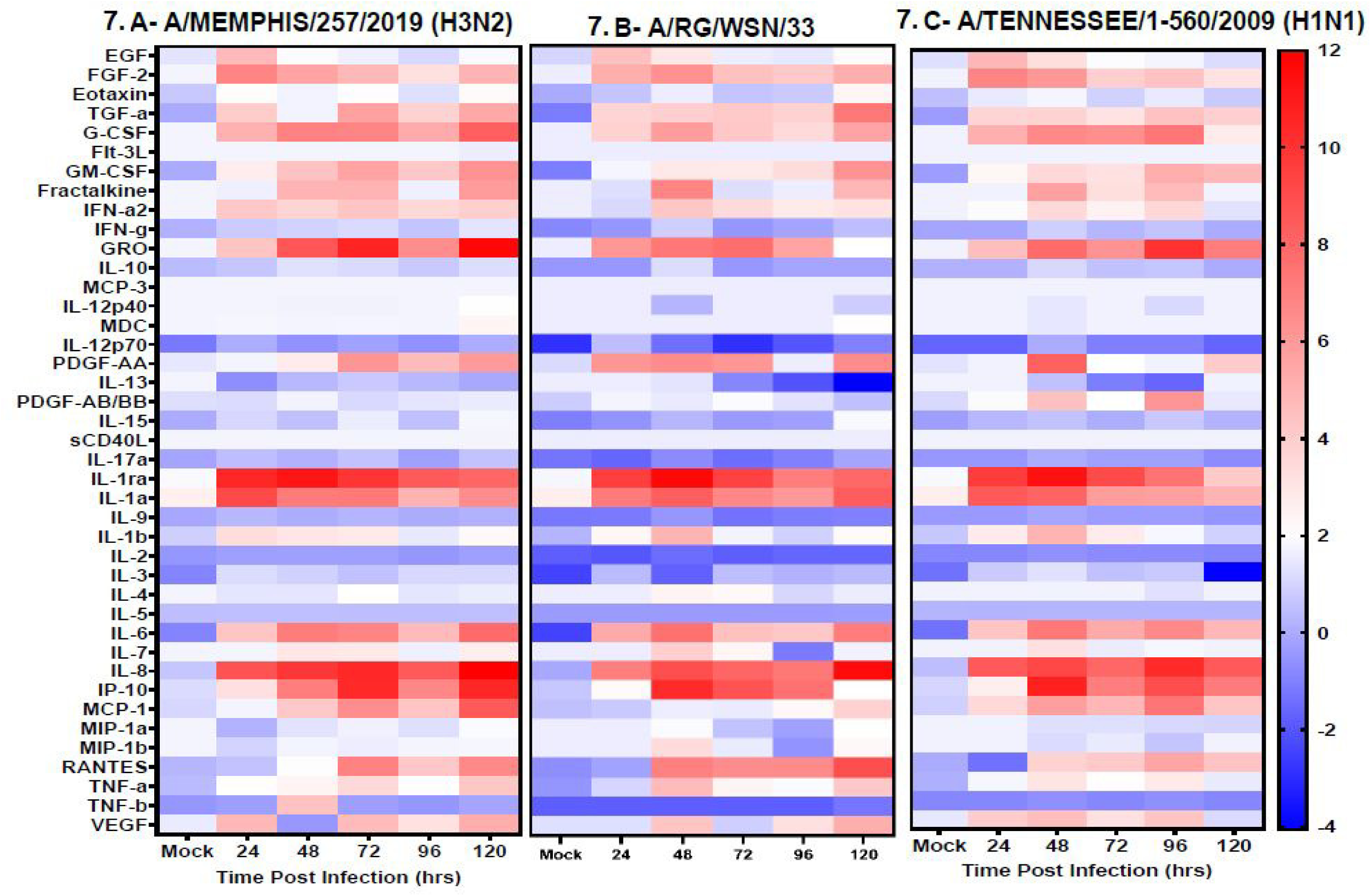
Heat map showing the cytokine immune response in HTECs at different times post infection. The values shown are fold changes from mock-infected cells. (A) H3N2. (B) pH1N1.

**FIG 8.**
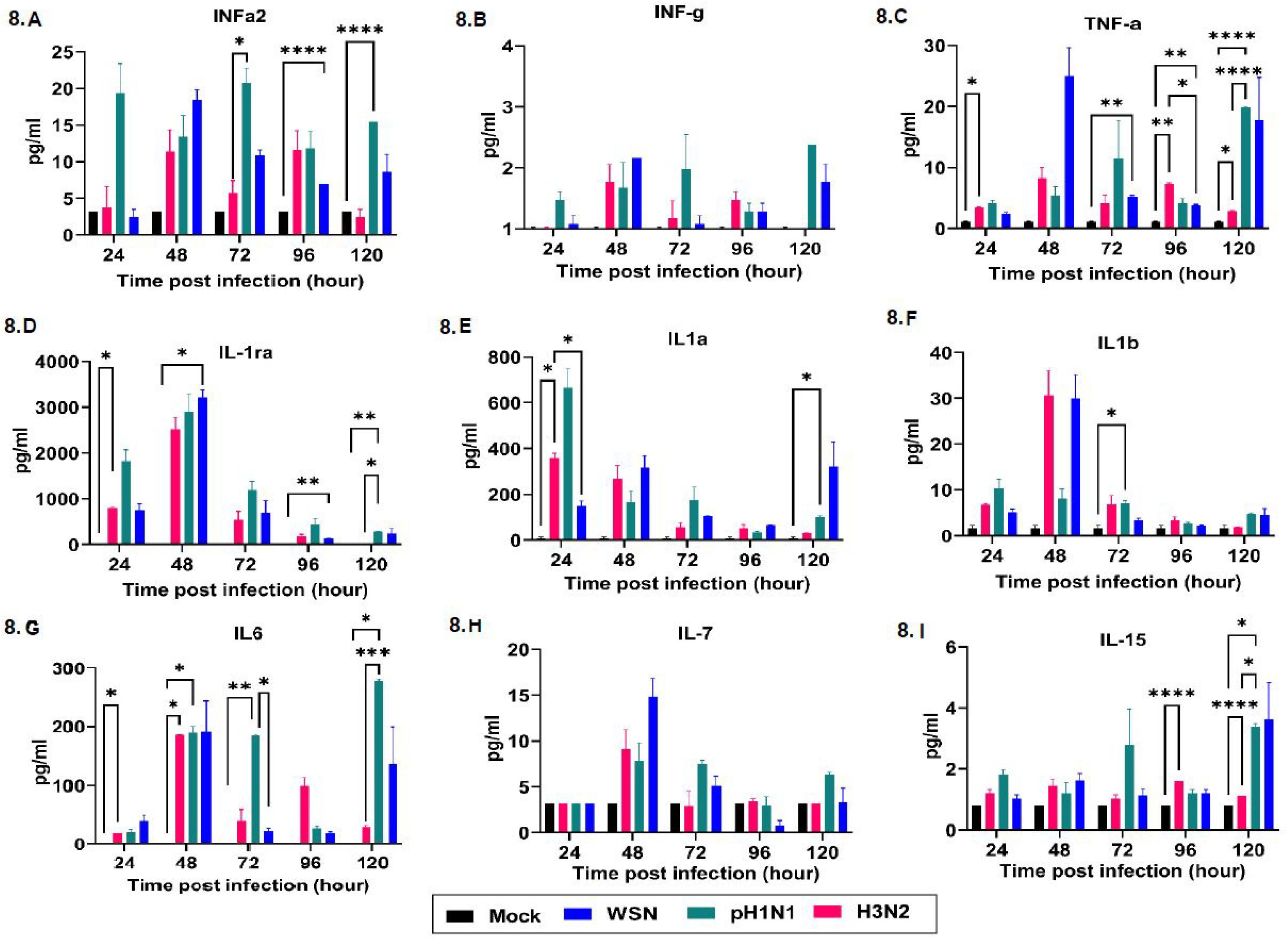
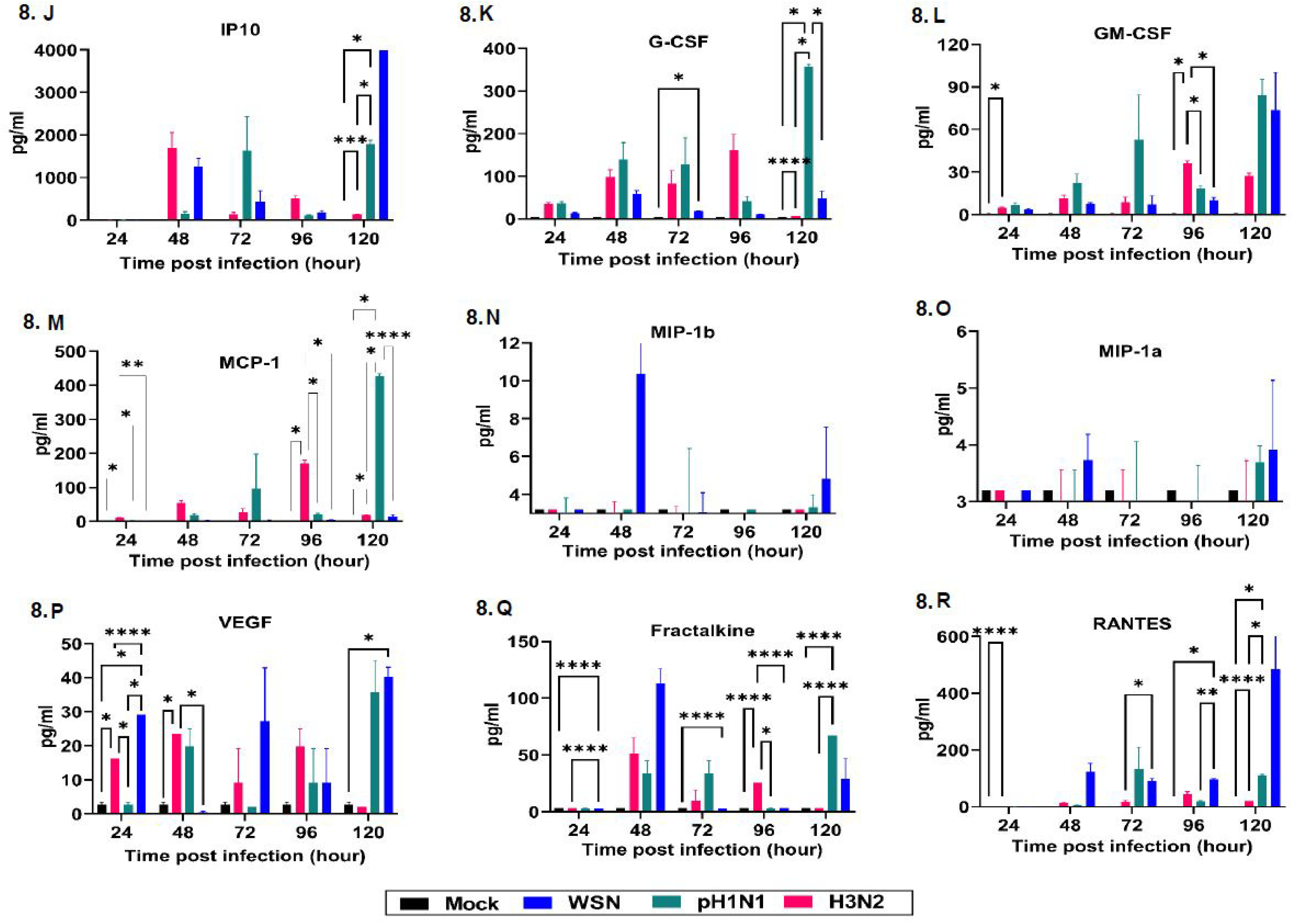
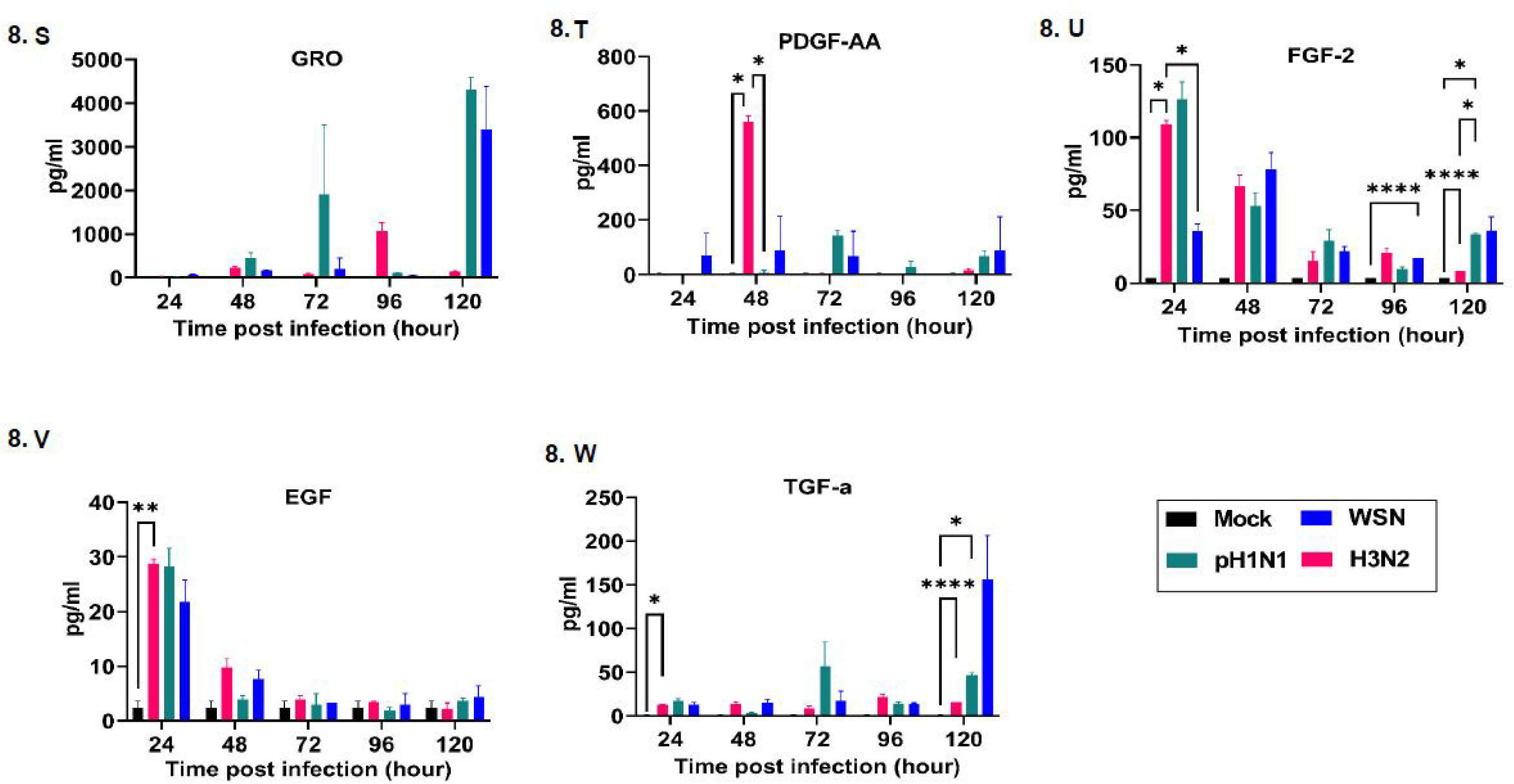
Cytokine secretion from HTECs upon infection with pH1N1, H3N2, and WSN IVs. The mean cytokine concentrations (pg/mL, log) detected at the indicated time points and infection conditions are shown (*n* = 5 replicates). Statistical significance was determined by two-way ANOVA and Tukey posthoc analysis, comparing each set of results to those of the mock virus. \**P <* 0.05; \*\**P <* 0.01; \*\*\**P <* 0.001; \*\*\*\**P <* 0.0001.

## DISCUSSION

Cell cultures of HTECs yielded heterogeneous squamous epithelial cells with clear differentiation and keratin expression patterns that are consistent with those of previous studies (8, 36). By using a panel of antibodies specific to different keratins, we confirmed that the tonsil epithelium consists of both stratified surface and reticulated crypt epithelial cells. *In situ* staining of the tonsil epithelium suggests that expression of the simple epithelial keratins K18 and K8, as well as K19, are specific to the crypt epithelial cells (5). K8 conventionally partners with K18 and was weakly expressed in keratinocytes of the surface tonsil epithelium but was strongly expressed in the reticulated crypts, specifically covering the upper layers of the crypts. Another cytokeratin that distinguishes the tonsillar crypts from surface epithelia is K19, which was variably expressed in the HTECs. K5 and K14 were strongly expressed in the surface epithelium. This suggests that our HTECs were highly differentiated and produced heterogeneous squamous cells, consisting of both surface stratified and crypt epithelial cells, consistent with those found in tonsil epithelial cells *in situ* (8, 37, 38). The stratified surface and reticulated epithelia cell populations within the HTECs contained large cytoplasm-to-nuclei ratios and heterogeneous squamous epithelial cells, further recapitulating human tonsil epithelial cells *in vivo*, with subpopulations of non-ciliated cells, few ciliated cells, and specialized cells with secretory functions.

Acquiring mutations and switching receptor-binding specificity between avian and human SA receptor preferences are key for IVs to transmit efficiently in humans and most likely occurs when adapting to new hosts (39, 40). We found that human-like *α*2,6-linked SA receptors predominated in the ciliated cells, and surface stratified epithelial cells contained abundant avian-like *α*2,3-linked SA receptors, which were more abundant and present on microvilli-containing reticular epithelial cells. Both *α*2,6-linked and *α*2,3-linked SA receptors were heterogeneously distributed. The HTEC surface and crypts were lined with pseudostratified columnar ciliated cells possessing both *α*2,6-linked and *α*2,3-linked SA receptors that were interrupted by patches of reticular epithelial cells.

IVs are transmitted by respiratory routes in humans and by the fecal–oral route in avian species. Human tonsils are located in the entrance of both the respiratory and digestive tracts, whereas avian tonsils are located only in the digestive tract (41). A typical feature of the tonsils is lymphocytic infiltration into the squamous epithelium of the crypts on its concave surface and the mucosal surface (42). We found the HTECs were permissive for the growth of both influenza A and B viruses. Heterogeneous variation occurred in the levels of immune mediators and cell death. A subset of cells, primarily ciliated cells, underwent apoptosis, but others, including non-ciliated cells, remained intact, despite being positive for IV nucleoprotein. Interestingly, colocalization occurred primarily with H3N2 in non-ciliated cells, whereas H1N1 mostly associated with ciliated cells.

Human innate and adaptive immune responses are activated shortly after IV infection (43). However, excessive or unbalanced immune responses can cause overproduction of both cytokines and chemokines, leading to severe inflammation and involving excessive recruitment of neutrophils and mononuclear cells at the site of infection, contributing to serious illness and mortality (44, 45). Many activities of innate and adaptive immune cells are coordinated by cytokines (44, 46–48). IL-6, IL-1α, IL-1β, IL-8, GM-CSF, and TNF-α are linked to protective Th1 responses (44, 49), effecting chemotaxis and activation of macrophages, neutrophils, and lymphocytes *in vivo*. The upregulation of these cytokines indicates that HTECs are an important source of proinflammatory cytokines in response to IV infection (48, 50). The tonsils provide defense against pathogens in the nasopharynx by infiltration of B and T lymphocytes into the heterogenous squamous epithelium of the crypts and the germinal center, in which B-memory cells are created and IgA secretory antibodies are produced (6). The germinal center reaction is critical for producing high-affinity and durable B-cell responses. Live attenuated IV vaccination provokes a germinal center reaction, which attracts B-cell clones, expanding the scope of protective antibodies caused by vaccinations (51). The excessive cytokine and chemokine response we observed in the HTECs indicates that the tonsils exacerbate virus-induced inflammation and pathology. Cytokine storm is associated with severe clinical outcomes during the course of IV infection (52–56). Several cytokines and chemokines, including VEGF, RANTES, MCP-1, IP-10, IL-8, IL-6, IL-1*α*, IL-1*β*, GRO, INF-*α*2, fractalkine, GM-CSF, TGF-*α*, FGF-2, and EGF, were secreted from HTECs at significant concentrations, consistent with dynamic communication between the tonsils and the immune system response earlier with H3N2 and later with pH1N1. High levels of IL-6 and IP-10 are linked to severe symptoms (57). The release of IP-10 from HTECs is also an important hallmark of seasonal IV infection, which can help clinicians make timely treatment decisions for patients with severe disease. Taken together, our findings demonstrate that several important cytokines are secreted from infected HTECs and that these cells are most likely to be the source of classic proinflammatory cytokines, such as IL-6 and TNF-*α*, which are elevated during systemic IV infections. The elicit production of the proinflammatory cytokines IL-15 and IL-6 in HTECs, suggesting a key role for these interleukins in the pathogenesis of epithelial cell damage and sore throat inflammation (58). Therefore, the cytokines release in relation to clinical findings, disease severity, and outcomes of children with IVs infection are needed. Our finding highlights the role of tonsils as a source of the previously reported proinflammatory cytokines, TNF, IL-6, IFN-*γ*1, IFN-*β*, IL-10, IL-8, and CXCL10, induced upon exposure to H3N2 (59). IV induction of IL-1*β* increases inflammation in lung cells; conversely, blockade of IL-1*β* signals with an IL-1*β* receptor antagonist or a neutralizing antibody alleviates inflammation (60), suggesting that IL-1*β* secretion by HTECs contributes to IV-induced inflammation. Thus, blockade of IL-1*β* signals can be a potential early therapeutic target for IV-induced inflammation (61). While IFN-*γ* is a particularly important factor in antiviral responses, and lower levels of IFN-*γ* confer weak antiviral ability (60), the lower levels of TNF-*α* and IFN-*γ* in our HTECs model may indicate weak anti-infection ability and the severe flu symptoms in children (62). We noted that HTECs upregulate VEGF in response to IV infection. This suggests that VEGF may be an early marker of IV infection and that HTECs are an important source of VEGF. MCP-1 levels were especially higher in patients with severe pneumonia who developed respiratory failure. These findings suggest that MCP-1 released by infected HTECs may contribute to the pathogenesis and severity of respiratory complications during IVs infection (63). While MIP-1β and IL-10 production is higher in fatal IV cases than in nonfatal cases (64). We did not observe any significant changes in MIP-1β in our HTEC model.

Our study reveals novel insights into the interplay between IVs and host signaling pathways in an underappreciated tissue such as the tonsils. The early release of cytokines and chemokines upon IV infection indicates that the tonsils are an important primary site of IV infections. Whether tonsillectomy affects the response to IV infections or vaccination is currently unknown. Tonsillectomy is a common surgical procedure in children (65), when the immune system is critically developing (66–69). We found a significant association in both immunity and cytokine storm in the HTECs. However, whether IV causes severe disease in individuals with or without tonsils and the prevalence of IV infections in both subpopulations remain to be determined. IV infections are well characterized to cause diverse outcomes that can result in severe complications and high mortality in some populations and mild symptoms in others. Thus, elucidating the factors related to these diverse outcomes is sorely needed.

In conclusion, the well-differentiated HTEC culture system we generated provides a valuable *in vitro* model for studying the cellular tropism and infectivity of IVs in human tonsils and may be valuable for developing an effective universal vaccine and therapies against different strains of IVs. Our results implicate the human tonsillar crypt epithelium as a site for IV replication that is permissive for the growth of different IV A and B strains.

## MATERIAL AND METHODS

### An air–liquid interface culture system for differentiated primary human tonsil epithelial cells

Primary HTECs were purchased from ScienCell Research Laboratories and were seeded on type I collagen (Sigma)–coated flasks supplemented with modified bronchial epithelial cell serum-free growth medium (70). The cells were transferred to type IV collagen-coated Transwell polycarbonate membranes (24-well, 0.4 *μ*m pore size, Corning Costar) in a cell density of 2.5 × 10^5^ cells/filter, with the air–liquid interface (ALI) medium supplemented with additives, as previously described (71). After the HTECs reached complete confluence, the cells were maintained under ALI conditions for 4–5 weeks at 37°C in a humidified 5% CO_2_ incubator.

### Morphological characterization and lectin staining of HTECs

Slides containing the well-differentiated HTEC cultures were treated the same way in all staining procedures. For indirect immunofluorescence analysis, transwell membranes containing HTECs were fixed and permeabilized for 10 to 15 min with 100% chilled acetone and then blocked with 5% bovine serum albumin. Primary antibodies against K5, K14, K8/K18, K19, villin (Abcam), and *β*-tubulin (Invitrogen) were typically diluted 1:200 and incubated for 1 h at room temperature. All of the antibodies were diluted in 2% bovine serum albumin and incubated at room temperature for 1 h. The slides were then washed three times with phosphate-buffered saline (PBS) and incubated with corresponding secondary antibodies. Secondary antibodies comprised goat anti-mouse or anti-rabbit antibodies conjugated to Alexa 488, Alexa 594, or Alexa 633 (1:5,000, Life Technologies). The cells were washed with PBS three times and visualized with a fluorescence microscope. Lectin staining was performed by using SNA or MAAII (Vector Laboratories) (72). Biotinylated lectin binding was visualized by fluorescence and confocal laser microscopy of streptavidin-conjugated Daylight 488 (Vector Laboratories). Nuclei were stained with 4′,6-diamidino-2-phenylindole and were embedded with ProLong Gold mounting medium (Life Technologies). ImageJ/Fiji software (National Institutes of Health) was used with the maximum intensity projection filter to process and analyze the confocal images. All experiments were repeated three times, with six fields per culture.

### Scanning electron microscopy

Scanning electron microscopy was performed as previously described (73). Briefly, cultured cells on membranes were fixed in 2.5% glutaraldehyde for 24 h and then treated with 1% osmium tetroxide for 2 h. This was followed by dehydration in an ascending series of ethanol and then dried with an E 3000 device (Polaron), adhered to stubs, sputter-coated, and examined under a scanning electron microscope (DSM940, Zeiss).

### Susceptibility of HTECs to IVs

Primary HTECs were grown at the ALI and infected apically with different IVs at various MOIs (Table 1). Three human H3N2 viruses (A/SWITZERLAND/8060/2017, A/HONG KONG/4801/2014, and A/MEMPHIS/257/2019), one swine H3N2 virus (A/SWINE/OH/16TOSU4783/2016), two human H1N1 viruses (A/TENNESSEE/1-560/2009 and A/BRISBANE/59/2007), two swine H1N1 viruses (A/SWINE/MN/37866/1999 and A/SWINE/NC/18161/2002), one swine H1N2 (A/SWINE/OH/09SW1486/2009), one avian virus (RGA/VIETNAM/1203/04), one influenza C virus (C/JOHANNESBURG/1/1966), and two influenza B viruses (egg-propagated B/BRISBANE/60/08 and cell-propagated B/BRISBANE/60/08) were used. We used A/RG/WSN/33 as a control virus because of its highly tropic and pathogenic effects.

### Viral replication kinetic in the well differentiated HTECs

To measure viral growth kinetics, three IV subtypes representing pH1N1 (A/TENNESSEE/1-560/2009), H3N2 (A/MEMPHIS/257/2019), and A/RG/WSN/33 were used. Well-differentiated HTECs (approximately 6 × 10^5^ cells/filter) were washed three times with PBS and inoculated with IVs from the apical side at a MOI of 0.1 for multiple-step kinetics and an MOI of 1 for single-step kinetics. After 2 h of incubation at 37°C, HTECs were rinsed with PBS three times to remove unbound viral particles, and fresh ALI medium was added. Infected HTECs were maintained under ALI conditions at 37°C in 5% CO_2_. At different time points, 100 *μ*L of DMEM was added to the apical surface, and the cultures were incubated for 30 min at 37°C. The harvested cells were collected at different times post infection. The infectivity of the viruses was evaluated by titrating the viruses by plaque assay or 50% tissue culture infectious dose.

### Immunofluorescence analysis for colocalization of viral protein

To determine the colocalization of viral proteins in HTECs, whole-filter cell cultures were fixed in 100% chilled acetone, blocked with 5% acetone, and incubated with primary monoclonal antibodies against influenza nucleoprotein (1:5,000) at room temperature for 30 min. The cells were then washed three times in PBS and stained with secondary goat anti-mouse Alexa 488 or Alexa 594 antibodies (1:5,000) (Molecular Probes) for 1 h in the dark. The cells were washed with PBS three times and examined with a fluorescence microscope (Nikon E400) or confocal laser microscope (Life Science Microscope, Keyence Corporation of America).

### Cytokine quantification

Multiplexed cytokine assays were performed according to the manufacturer instructions by using a Bio-Plex Pro Human cytokine screening panel (Biorad) containing 48 human cytokines, and the spectral intensities were quantified with a Luminex instrument. Cytokine concentrations were calculated by interpolating values from a standard curve via 5PL curve fitting. Samples with values below the detection limit were assigned a value of 1 to enable the use of a log scale for calculating fold changes and graphing. IL-11 was quantified by using a Human IL-11 Quantikine ELISA Kit according to manufacturer instructions (R&D Systems). Statistical analysis was performed on log_10_-transformed values with two-way ANOVA tests and Tukey correction. To identify the cytokines secreted by uninfected HTECs, the following criteria were used: < 10-fold increased cytokine signal at different time points compared to media. To identify significantly increased cytokine concentrations upon infection, an adjusted *P* value (< 0.01) between the cytokine signals at different time points *vs.* those of uninfected controls was used.

### Statistical analyses

All data shown are means ± standard error. All statistical analyses were performed with Prism 9 software (GraphPad Software).

## Acknowledgments

We are grateful to the imaging help provided by Ryan Kelly, from the Keyence Corporation of America. We are thankful for Bindumadhav Marathe and John Franks for their technical help, for Dr. Cam Robinson for his help in scanning microscopy, and for the excellent scientific editing of the manuscript by Nisha Badders.

## Author Contributions

F. O. performed the experiments, analyzed the data, and wrote the manuscript; A.S. analyzed the data; R. W. supervised the study; R.J.W. supervised the study, analyzed the data, Funding, and wrote the manuscript. All authors have read and agreed to the published version of the manuscript.

